# Efficient multi-gene expression in cell-free droplet microreactors

**DOI:** 10.1101/2021.11.10.468096

**Authors:** Ana Maria Restrepo Sierra, Stefan T. Arold, Raik Grünberg

## Abstract

Cell-free transcription and translation systems promise to accelerate and simplify the engineering of proteins, biological circuits and metabolic pathways. Their encapsulation on microfluidic platforms can generate millions of cell-free reactions in picoliter volume droplets. However, current methods struggle to create DNA diversity between droplets while also reaching sufficient protein expression levels. In particular, efficient multi-gene expression has remained elusive. We here demonstrate that co-encapsulation of DNA-coated beads with a defined cell-free system allows high protein expression while also supporting genetic diversity between individual droplets. We optimize DNA loading on commercially available microbeads through direct binding as well as through the sequential coupling of up to three genes via a solid-phase Golden Gate assembly or BxB1 integrase-based recombineering. Encapsulation with an off-the-shelf microfluidics device allows for single or multiple protein expression from a single DNA-coated bead per 14 pL droplet. We envision that this approach will help to scale up and parallelize the rapid prototyping of more complex biological systems.

## 1 Introduction

Cell-free extracts played an instrumental role in early biochemical research [1, 2]. More recently, cell-free protein expression systems were re-discovered [3] and optimized as versatile platforms for the prototyping of synthetic genetic circuits [4], engineered proteins [5] or multi-enzyme biosynthesis pathways [6]. For bioengineering purposes, the main advantages of cell-free expression systems lie in their flexibility, speed and reduced complexity [7]. Synthetic protein-coding DNA can be applied directly to the reaction, without previous sub-cloning into plasmids or genomes [8, 9]. Expression products are available within hours, and can then be characterized within the same reaction without cell lysis or lengthy extraction procedures [10]. Expression conditions can be manipulated beyond what is possible in the tightly regulated intracellular environment. Moreover non-natural molecules or even toxic and membrane disrupting molecules can be produced [11, 12, 13, 14, 15, 16]. The latest generation of cell-free reactions are re-constituted from a limited set of purified components [17]. These defined systems further lower reaction complexity, facilitate product identification, improve reproducibility and minimize side-reactions.

However, engineering approaches that rely on large-scale randomization and selection are still bound to cells. Without compartmentalization, cell-free systems that are testing different genetic programs have to be physically divided and evaluated one by one. In 1998 Tawfik et al. started addressing this limitation by encapsulating cell-free reactions in water-in-oil emulsions [18]. Cell-free droplet production was later streamlined by the introduction of custom microfluidic setups [19, 20]. Current microfluidic approaches indeed improve throughput and give better control over droplet size and composition. Limiting dilution of DNA was then used to create “clonal” droplets with only one DNA molecule (encoding GFP) per compartment [21].However, protein expression levels were low, which was not surprising as cell-free reactions are typically not efficient with low quantities of template DNA [22]. Several studies thus introduced a DNA amplification step prior to in-droplet cell-free expression [23, 24, 25, 26]. However, this approach implies droplet merging or micro-injection steps, which rely on custom microfluidic designs that can be tedious to implement. More recently, micro-beads have been explored as DNA-binding scaffolds for cell-free reactions, and it was shown that proteins can be expressed in larger microliter-scale reactions directly from plasmid DNA bound to beads [27]. Independently, novel gene synthesis methods have been proposed to assemble libraries of a short gene from oligonucleotides within emulsion droplets [28, 29]. A recent study then combined similar technology with cell-free expression for the *in-vitro* directed evolution of an affibody [30]. So far, these studies have focused on single proteins. Combinatorial engineering of more complex systems such as pathways or genetic circuits [9, 31] remains out of reach as current methods do not support the introduction of genetic diversity while also ensuring high co-expression levels of multiple proteins within cell-free micro-reactors.

We here describe a proof-of-concept *in-vitro* bioengineering platform for the rapid cell-free expression of multiple proteins in microfluidic droplets. We co-encapsulate cell-free reactions with DNA-coated micro-beads to generate droplets with high concentration of template DNA that can nevertheless differ between the drops. We optimized the loading of template DNA onto these beads. We then evaluate a step-wise solid-phase DNA assembly which simplifies multi-protein expression in droplets without the need of cloning. Our approach utilizes readily available off-the-shelf hardware and reagents which makes it accessible for researchers who are not microfluidics specialists. We anticipate that the co-encapsulation of multiscistronic DNA-coated beads with cell-free expression systems will enable rapid and high-throughput engineering of genetic circuits and metabolic pathways.

## 2 Materials and Methods

### 2.1 DNA constructs

The DNA constructs used in this study are described in Supporting Information, Table S1. Annotated DNA sequences are available online: [https://github.com/strubelab/dropletXpress]. Protein-coding DNA sequences were codon-optimized with the COOL web server [32] to match highly expressed *E. coli* genes in terms of (i) individual codon-usage, (ii) codon-pair usage, and further optimized for (iii) hidden stop codons, (iv) ≈50% GC content, (v) removal of consecutive nucleotide repeats >4 bp, (vi) removal of any sequence repeats ≥ 8 bp. Linear dsDNA fragments were synthesized by Integrated DNA Technology (IDT) and cloned via isothermal assembly [33] into our in-house pJEx411c vector backbone (KanR, lacI, lacO, T7p), an expression vector modified from the original DNA 2.0 pJExpress411 to include a short leader peptide expression cassette out of frame, which insulates the RBS from expression inhibition by mRNA secondary structure [34]. The Bxb1 expression construct (sb0201) was gene-synthesized by Twist Bioscience (US).

### 2.2 dsDNA Fragments for binding and assembly on beads

The sequences of oligonucleotides used in this study are listed in Supporting Information, Table S2. All oligonucleotides were synthesized by IDT (US). Short dsDNA fragments for binding on beads were created by annealing the complementary oligonucleotides aro0024 and aro0025 in a 50 *µ*l reaction containing 10 *µ*l of 100 *µ*M of each primer, 5 *µ*l of 10X NEB2 buffer (NEB, US), and 35 *µ*l of nuclease-free water. The reaction was heated to 95 °C for 2 min on a thermocycler and gradually cooled down to 25 °Cover 45 min. All PCR products utilized in this study are listed in Table S3 in Supplemental Materials, along with templates and primers used for their construction. Reactions were performed with Phusion High-Fidelity Master Mix and GC buffer(NEB, US) using 1 to 30 ng of plasmid DNA templates. Thermocycler protocols are specified in suppplemental tables S4, S5, and S6.

### 2.3 Direct DNA binding on beads

Polystyrene microspheres coated with Streptavidin (mean diameter 4.95 µm) were purchased from Bangs Laboratories (US). Washes and incubations were preferably performed in low-binding tubes (Eppendorf, Germany). Beads were washed and centrifuged three times for 3 min at 21,000 g in 100 *µ*l of binding buffer (10 mM Tris-HCl, 1 M NaCl, 1 mM EDTA, 0.0005 % TritonX-100). Beads were then resuspended in 10-50 *µ*l of binding buffer containing 5 to 300 pmol of template DNA. Reactions were incubated at room temperature for 3 h or overnight under continuous mixing using a MonoShake Microplate Mixer (Variomag) at maximum speed (2000 rpm). Following binding, beads were centrifuged for 3 min at 21,000 g and the supernatant with unbound DNA was removed. The amount of DNA bound was estimated by measuring the DNA concentration in the supernatant using a Nanodrop photospectrometer (ThermoFisher, US). Beads were repeatedly washed with binding buffer and nuclease-free water until the DNA concentration in the supernatant was close to zero. DNA-immobilization on magnetic beads (Dynabeads M-280, mean diameter 2.8 µm, Thermofisher, US) was performed following the same protocol.

### 2.4 Protein expression and purification

Expression plasmids were transformed into *E. coli* BL21(DE3). Starter cultures (2xYT, 50 *µ*g/ml kanamycin) were inoculated from single colonies, grown over night at 37 °C and then used for 1:100 inoculation of 1 l production cultures (2xTY, 50 *µ*g/ml kanamycin, 1% glucose). Production cultures were grown shaking to until an O.D. at 600nm of 0.6 - 0.8, induced with 0.5 mM IPTG (final concentration) and incubated over night at 20 °C. Cells were harvested by centrifugation for 15 min at 6000 g and 4 °C, washed once in 15 ml PBS, weighed and stored at -80 °C. Pellets were resuspended in 5 ml per g of pellet with a lysis buffer consisting of 100 mM Tris-HCl (pH 8.0), BugBuster concentrate (Millipore),150 mM NaCl, 1 mM EDTA, 5% glycerol, 1 mM TCEP, freshly supplemented with protease inhibitor cocktail (Thermofisher) at 1 tablet/50 ml, 350 U/ml Benzonase (Millipore) and 0.5 mg/ml lysozyme (Sigma). The lysis mix was incubated for 45 min rotating at 4 °C. Cell debris and insoluble protein was removed by 45 min centrifugation at 80,000 g and 4 °C. The supernatant was filtered over Miracloth (Millipore) and on one 5 ml StrepTrap HP column (GE Healthcare, US) equilibrated in binding buffer (100 mM Tris-HCl pH 8.0, 150 mM NaCl, 1 mM EDTA, 5% glycerol, 0.5 mM TCEP), washed with 5 column volumes binding buffer and then eluted with 2.5 mM Desthiobiotin (ThermoFisher, US) in binding buffer on an Äkta Pure FPLC (GE healthcare) with sample pump. Fractions were pooled, concentrated in Amicon-Ultra 10 kD spin concentrators (Millipore) and subjected to size exclusion chromatography on a Superdex 75 16/600 column (GE Healthcare) equilibrated in 20 mM HEPES pH 7.4, 300 mM NaCl, 10% glycerol, 50 *µ*M EDTA, 0.5 mM TCEP. Protein fractions were pooled, concentrated, aliquots flash-frozen in liquid nitrogen and stored at -80 °C.

Cell pellet with Bxb1 was lysed in 25 mM Tris-HCl (pH 7.4), 0.5 M NaCl, 40 mM Imidazole, 10% Glycerol, 1 tablet/50 ml COmplete Protease Inhibitor (EDTA-free, Sigma, US), 1 mM TCEP, Bugbuster, lysozyme, Benzonase (25 U/ml) as above. Bxb1 recombinase was captured on a 5 ml HisTrap HP column (GE, US) in standard His binding buffer (25 mM Tris-HCl pH 7.4, 0.5 M NaCl, 40 mM Imidazole, 10% Glycerol, 1 mM TCEP), eluted with a gradient into a higher salt elution buffer (20 mM Tris pH 8.0, 1 M NaCl, 0.5 M Imidazole, 2 mM EDTA, 1 mM TCEP) and dialyzed overnight into 20 mM Tris-HCl pH 8.0, 2 mM EDTA, 1 mM DTT, 1 M NaCl and concentrated. For storage at -80 °C, glycerol was added for a final composition of 8.5 mM Tris-HCl, 0.9 mM EDTA, 0.4 mM DTT, 425.5 mM NaCl, 50% glycerol. The product of this single-step purification was more than 95% pure (Figure S3) and directly used for coupling reactions. All proteins expressed in this study, either in PURExpress cell-free reactions or in *E. coli* BL21(DE3), are listed in Table S7.

### 2.5 On-bead DNA concatenation with Golden Gate or Bxb1

For Golden Gate assembly or Bxb1 recombination, the first linear DNA fragment with either a BsaI restriction site or a Bxb1 recombination site at the 3’ terminal (following the T7 terminator), was bound to streptavidin beads as described above. DNA-coated beads were then resuspended in nuclease-free water. PCR-amplified linear DNA, with the complementary restriction or recombination site, was added at a molar ratio of 1:1 or 1:2. Golden Gate reactions were performed in 1X T4 ligase buffer (NEB) with 1000 units of T4 DNA Ligase (NEB), and 30 units of BsaI-HF V2 (NEB) in a total volume of 35 *µ*l per reaction. Reactions were cycled 30 times with 5 min incubation at 37 °C and 5 min incubation at 16 °C (no heat inactivation). Bxb1 recombinase reactions were performed in 10 mM Tris-HCl pH 8.0, 100 mM KCl, 10 mM EDTA in a total volume of 35 *µ*l per reaction. Purified Bxb1 was added at 55 fold molar excess over DNA bound on the beads (7.7 pmol Bxb1 per estimated 0.14 pmol of bound DNA). Bxb1 storage buffer distorted the final salt concentration but we ensured that it never exceeded 150 mM NaCl. Reactions were incubated for 3 h at 30 °C without heat inactivation step. After concatenation, beads were washed several times with binding-washing buffer (see above), and washed and then resuspended in nuclease-free water for another cycle of concatenation or, finally, for PUREexpress reactions. PCR fragments and corresponding primers are specified in Table S3.

### 2.6 Cell-free expression in bulk reactions

25 *µ*l reactions were set up following the PURExpress protocol (In Vitro Protein Synthesis Kit, NEB) in black low-volume 384-well polystyrene plates (Corning). Both linear DNA in solution and DNA-coated beads were used as templates for bulk protein expression at a final DNA concentration ranging from 10 ng/ *µ*l to 40 ng/ *µ*l. DNA-coated beads were resuspended in 7 *µ*l of nuclease-free water before being added to the cell-free reaction. Reactions in plates were covered with plastic seal and incubated for 6-8 hours at 37 °C. Protein expression was quantified by fluorescence measured on a Tecan M1000 plate reader with the following parameters: mNeonGreen (excitation 506 nm, emission 517 nm), LSSmOrange (excitation 437 nm, emission 572 nm), mScarlet-I (excitation 569 nm, emission 593 nm). Top and bottom fluorescence readings with a signal gain of 60, 80, and 100, were recorded from each well. Three calibration curves were prepared with seven concentrations of purified fluorescent proteins (Figure S1).

### 2.7 Cell-free expression in droplets from DNA in solution or on beads

Encapsulated protein expression both from DNA in solution and from DNA on beads was performed on a commercial *µ*Encapsulator system (Dolomite Microfluidics) holding a two-reservoir chip and a 30 µm fluorophilic droplet generation chip. The two-reservoir chip was filled with 30 to 80 *µ*l PURExpress in one reservoir, and the same volume of either DNA in solution or DNA-coated beads in the other. Both reservoirs were then filled up to their total capacity of 100 *µ*l with FC-40 oil. The two reagent streams were driven by a single Dolomite P-pump at a flow rate of 10 *µ*l /min under the control of a single 1-50 *µ*l /min flow sensor. This flow path was split by a T-junction to obtain 5 *µ*l /min flow for each reagent stream. The length of tubing after the T-junction was identical in order to ensure equal flow. A second Dolomite P-pump regulated by a 30-1000 *µ*l /min flow sensor provided carrier fluid (FC-40 with 2% PicoSurf filtered over 0.22 um) at 30 *µ*l /min. FEP tubing with 0.25 mm inner diameter and 0.8 mm (1/16”) outer diameter and flow regulation valves were used as recommended by the manufacturer. For experiments with DNA in solution, DNA was diluted in nuclease-free water to 20 ng/ *µ*l, corresponding to 10 ng/ *µ*l final concentration in droplets. Otherwise, DNA-coated beads were diluted in nuclease-free water with 18 % OptiPrep (Sigma-Aldrich) to a final concentration of 1 to 2 beads per 14 pL droplet. In order to avoid clogging, beads were vigorously vortexed before loading into the reservoir chip. If clogging nevertheless occurred during droplet generation, the *µ*Encapsulator was opened and the droplet generation chip was flushed through each channel using a 1 ml syringe with filtered Milli-Q water, dried with pressurized air, and reconnected. Droplets were collected in a 1.5 Eppendorf tube and incubated at 37 °C for 6-8 h.

### 2.8 Confocal microscopy

Single emulsion droplets were loaded into Hollow Rectangle Capillaries ID 0.05 × 0.50mm (CM Scientific) or into Countess Cell Counting Chamber Slides (ThermoFisher Scientific) for microscopy imaging. Images were taken on a CLSM LSM710 (Zeiss) upright confocal microscope with 10X objective. Parallel images were taken with transmitted light and the corresponding fluorescence laser and filter setting for mNeonGreen (excitation 488 nm, emission 515-564 nm) or LSSmOrange (excitation 440 nm, emission 536-706 nm) or mScarlet-I (excitation 561 nm, emission 589-740 nm). A droplet-based protein concentration calibration curve was prepared from purified mNeonGreen protein. Four different concentrations of mNeonGreen were encapsulated with the *µ*Encapsulator system and the fluorescence was measured under identical conditions (supplemental figure S2). Images were analyzed in MATLAB with a script modified from a script kindly provided by Daniela Garcia (UPF, Barcelona, Spain) ([https://github.com/strubelab/dropletXpress]) and protein concentrations were determined from the median fluorescence intensity.

## 3 Results

### 3.1 Expression of single and multiple proteins from DNA-coated beads

We first carried out microliter-scale cell-free reactions to compare protein expression from regular DNA in solution and DNA immobilized on beads. We loaded 500 ng of dsDNA onto 35 · 10^5^ of beads, as recommended by the manufacturer. We found that expression from DNA on beads gives protein levels comparable to expression from equivalent DNA amounts in solution (Figure 1a). We also observed that the position of the biotin attachment (3’UTR or 5’UTR), did not have an influence on protein expression levels (Figure 1a).

**Figure 1.**
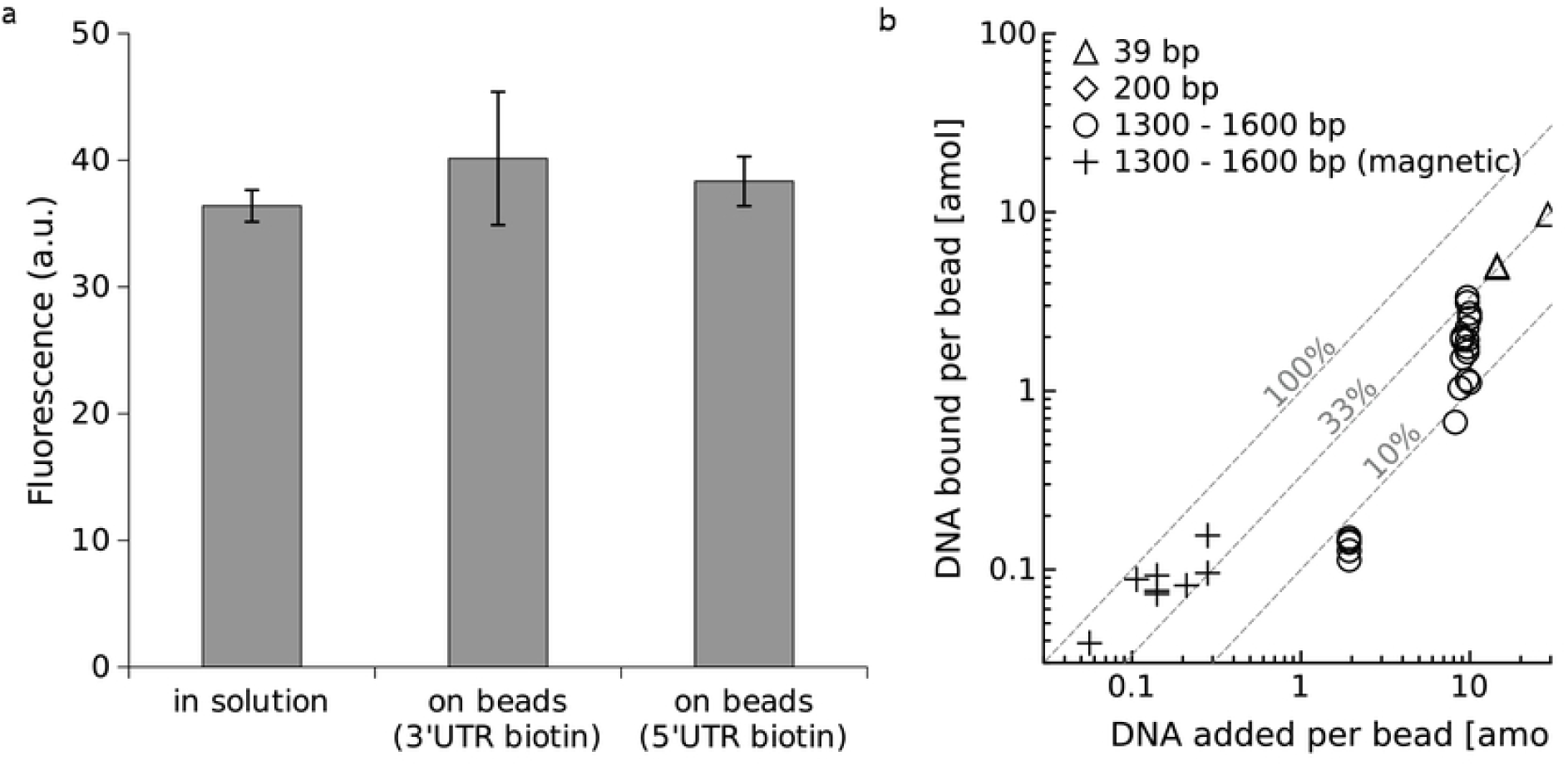
DNA immobilization on beads and expression in µl-scale reactions. (a) Comparison of fluorescent protein expressed in 25 *µ*l reactions from DNA in solution or equivalent amounts of DNA directly immobilized on beads. Error bars show standard deviation calculated from three replicates. (b) Optimization of direct DNA loading: biotinylated dsDNA fragments of various sizes were incubated with Streptavidin-coated micro beads. Dashed lines: DNA binding efficiency

However, under these conditions, the amount of DNA carried by a single bead into a 14 pL microfluidic droplet would translate to only about 5 ng/ *µ*l which is at the lower end of DNA concentration recommended by the manufacturer of the cell-free system. We achieved an average of 20-fold higher loading levels per bead by over-saturating them with input DNA at the cost of leaving more DNA unbound in solution (Figure 1b). There was thus a trade-off between overall binding efficiency (better at low concentrations) and the DNA quantity bound per bead which had been previously speculated on [35] (Figure S4). Maximum loading levels were reached with short (39 bp) double-stranded oligonucleotides as these could be provided at even higher starting concentrations (Figure 1b).

Rather than forcing the, likely, inefficient direct binding of much larger multiscistronic templates, we decided to test whether longer DNA constructs could be built on beads with solid-phase DNA assembly techniques. This approach could also potentially enable combinatorial multi-gene expression from individual single-gene DNA fragments without the need for tedious cloning steps. To evaluate this concept, we pre-loaded beads with short biotinylated double-stranded oligonucleotides which would then serve as specific anchors for longer DNA fragments in a solid-phase DNA assembly reaction. We tested two DNA assembly methods, Golden Gate [36] and Bxb1 recombineering [37, 38] for the assembly of, initially, only one protein-coding DNA construct onto these beads. We compared this assembly reaction with the direct binding of the equivalent biotinylated DNA fragment under efficient binding conditions (i.e. in the non-saturating DNA regime). We used these beads as a template for *µ*l -scale PURExpress cell-free reactions and quantified the concentrations of a fluorescent protein expressed from either the assembled or from the directly immobilized DNA (Figure S5). Both Golden Gate and Bxb1 DNA assembly strategies yielded protein expression levels comparable to the control reaction where the protein-coding DNA construct had been directly bound to the beads. Thus, both assembly method allow for the selective and efficient coupling of genes to beads.

We then used this strategy for a sequential solid-phase assembly of multicistronic (multi-gene) constructs. Micro-beads were first saturated with a biotinylated mScarlet-encoding DNA fragment to which two additional DNA fragments, encoding for mNeon and LSSmOrange, were sequentially assembled through either Golden Gate ligation or and Bxb1 recombineering. The beads from either approach were again subjected to *µ*l -scale PURExpress cell-free expression reactions. After calibration against purified proteins and thanks to minimally overlapping excitation and/or emission spectra, we could reliably deconvolute and quantify the concentrations of all three fluorescent proteins within the same reaction. Both methods demonstrated efficient solid-phase DNA assembly for multi-protein expression in the bulk cell-free reaction (Figure 2). Again, product concentrations were comparable between DNA provided in solution or concatenated on beads.

**Figure 2.**
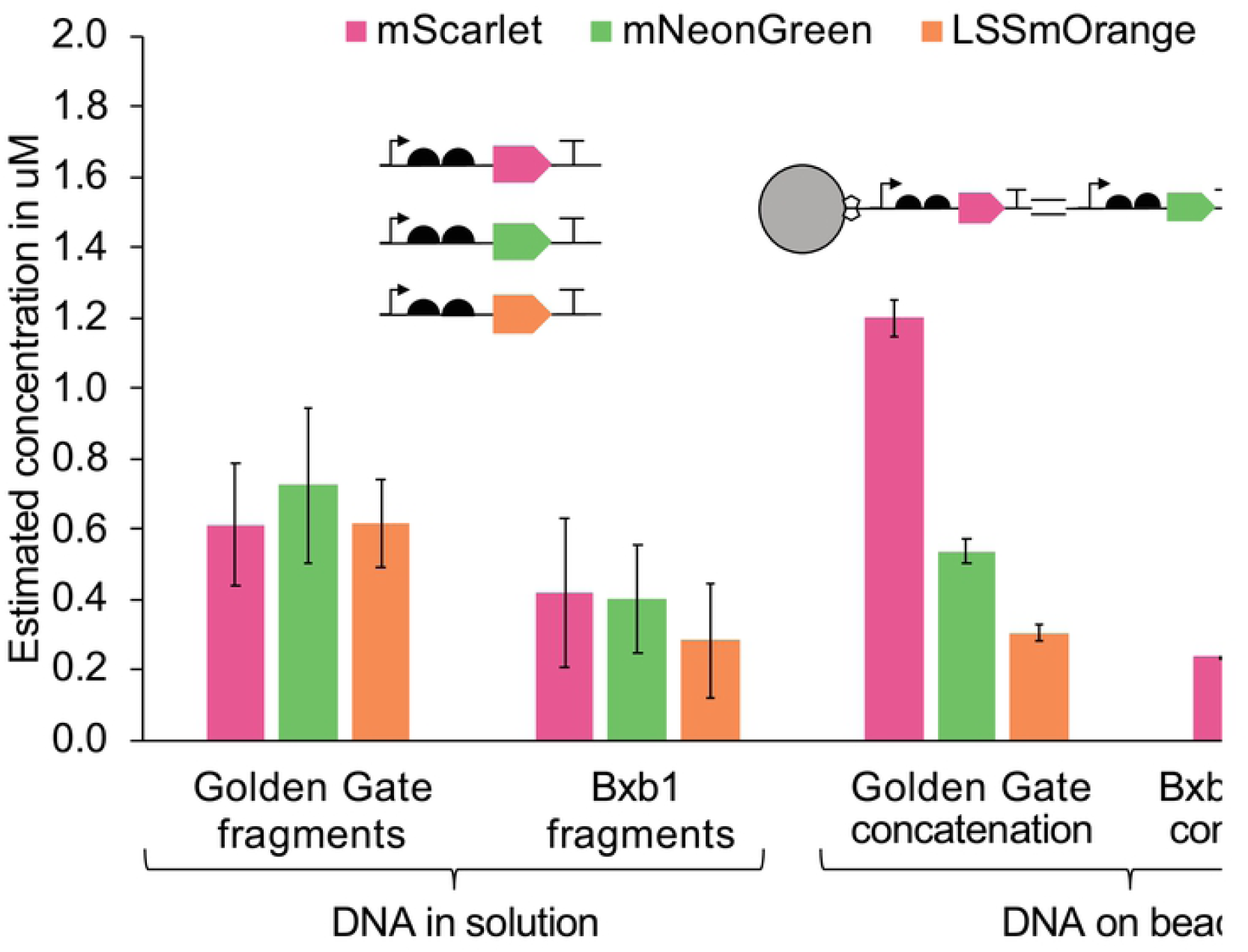
Co-expression of three proteins in bulk reactions (25 *µ*l) from linear DNA templates supplied in solution (control) or from DNA immobilized on micro-beads. Equal quantities of PCR fragments encoding red, green and orange fluorescent proteins were either directly mixed into PureExpress reactions (controls, left) or were sequentially immobilized on beads using two different solid-phase gene coupling strategies (Golden Gate and Bxb1 Integrase concatenation). Expression was measured by fluorescence and calibrated against purified fluorescent proteins. Error bars indicate the standard deviation among triplicates.

Judging from the control reactions, the addition of all three DNA fragments in 1:1:1 ratio led to similar protein expression levels. When using the Bxb1 integrase for DNA assembly, all three fluorescent proteins were evenly expressed, independently of their position in the assembled construct. Hence, we conclude that this assembly method had a coupling efficiency of close to 100%. By contrast, expression levels decreased from first to last fragment for beads loaded by Golden Gate assembly. Nevertheless, both methods allowed a robust and high protein co-expression. Although the Golden Gate assembly was somewhat less efficient, it is more accessible as the enzymes are commercially available. In summary, our experiments in bulk solution confirmed that one or multiple proteins can be efficiently expressed from DNA directly assembled on beads.

### 3.2 In-droplet protein expression

As a next step, we evaluated protein expression in microfluidic droplets, at first from a linear DNA fragment provided in solution and encoding the mNeonGreen fluorescent protein. We tested different flow rates and configurations for reconstituting cell-free reactions in a commercial two-reagent-stream droplet generation setup. We settled on the most reproducible set-up: an equal flow rate with one stream supplying DNA and the other providing the cell-free system (Figure 3a,b). We adjusted the in-droplet DNA concentration to 10 ng/ *µ*l, which is recommended by the manufacturer as starting DNA concentration in standard 25 *µ*l PURExpress reactions. In-droplet protein expression reached concentrations of 0.4 - 0.8 µM *µ*M (Figure 3c) and corresponds to approximately 15% of what we obtained in a standard 25 *µ*l reaction. We then immobilized the same template by saturation of beads with biotinylated DNA. Beads were encapsulated at approximately one bead per droplet resulting in a nominal DNA concentration of 120 ng/ *µ*l. In-droplet expression of mNeonGreen from beads reached protein concentrations between 0.2 - 0.5 *µ*M. Despite the higher DNA template concentration in single-bead droplets, protein expression yield was minimally lower than from droplets with DNA template in solution (Figure 3c). One explanation might be that template DNA became less available for transcription on the very densely loaded beads under our saturating conditions. We noticed that some droplets contained beads (single or multiple) but did not express any fluorescent protein. We speculate that clustering of beads [35] may sometimes prevent DNA capture. Furthermore, flow rate variability during droplet generation may perturb the concentration of cell-free reagents.

**Figure 3.**
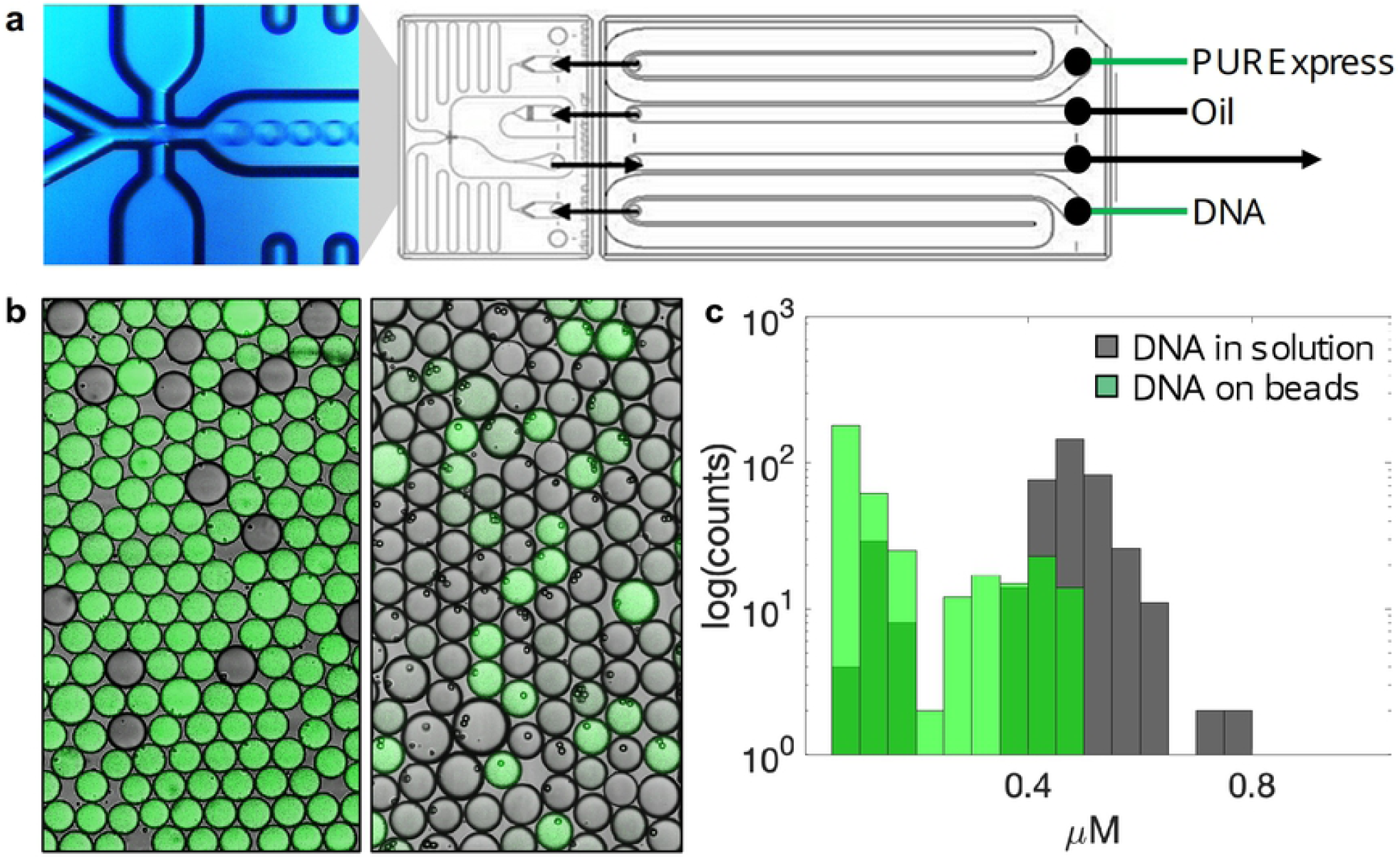
Droplet-based expression of mNeonGreen fluorescent protein from DNA in solution and from single DNA-coated beads. (a) Schematic microfluidics setup with the actual droplet generation junction shown on the left. (b) Fluorescent microscopy images of representative droplets with white light transmission channel overlay. Left: expression from DNA in solution, Right: expression from DNA-coated beads. (c) protein concentration distribution over 401 droplets (DNA in solution), and 354 (DNA on beads. One bead per droplet).

Encouraged by these results, we now went on to try creating genetic diversity between droplets. We bound biotinylated DNA coding for mNeonGreen, mScarlet-I, or LSSmOrange onto three separate batches of beads. After separate DNA-binding and washing steps, we mixed these beads and used them for a single-bead cell-free encapsulation experiment. As we expected, we saw a clear separation of color between different droplets (Figure 4a). This validates that each droplet utilized multiple copies of a single type of DNA template without noticeable cross-contamination between the droplets.

**Figure 4.**
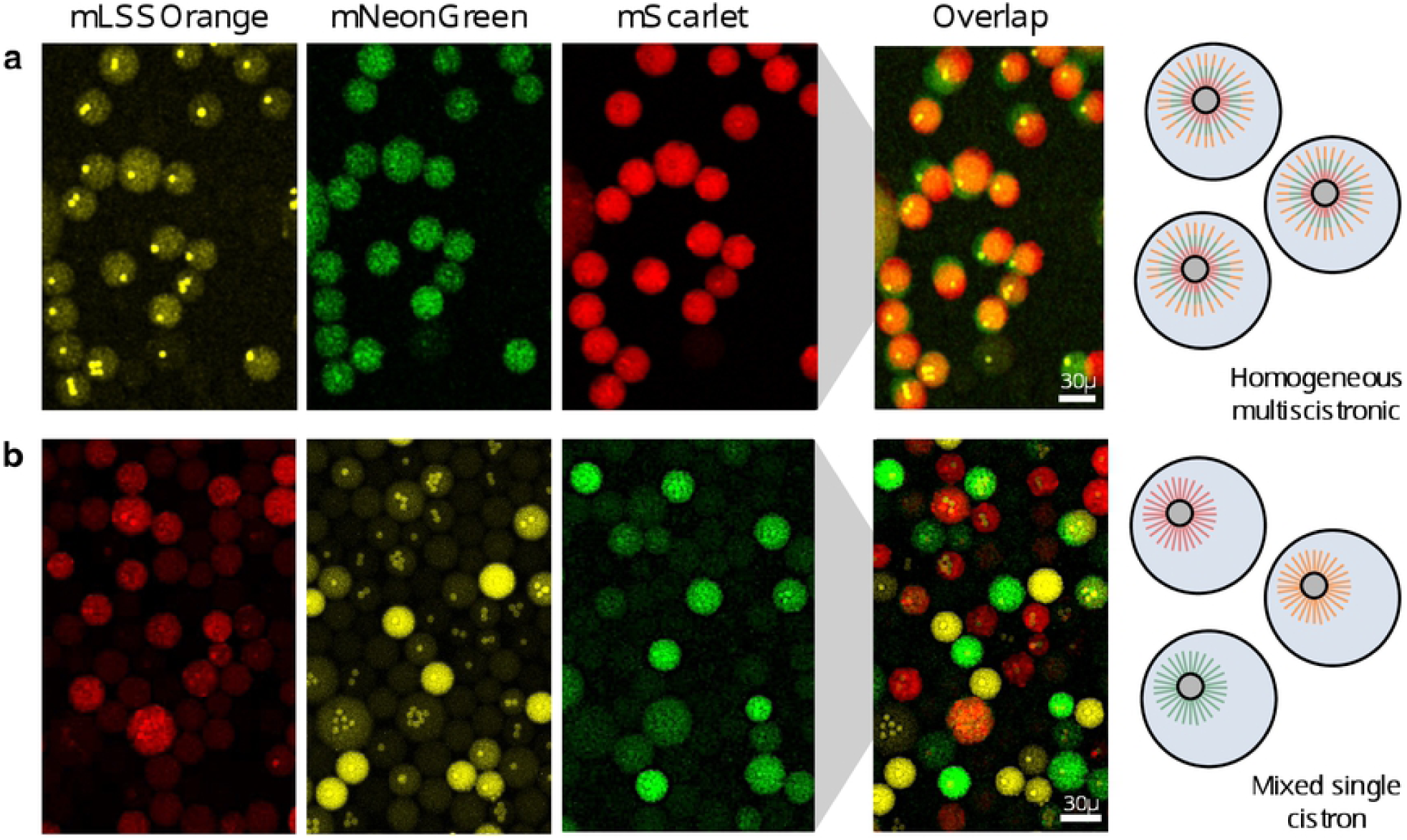
Co-expression or parallel expression of three different proteins from beads in droplets. (a) For co-expression in the same droplet, three genes encoding different fluorescent proteins were concatenated onto beads using Golden Gate assembly and beads were encapsulated with cell-free system. Co-expression of LSSmOrange, mNeonGreen, and mScarlet-I in all droplets is evident from imaging in the three different emission channels (note that droplets may be moving between images). (b) For parallel expression, genes were immobilized onto beads separately, beads were mixed and encapsulated together. Imaging in the three emission channels shows that most droplets only express one of the three fluorescent proteins.

Having separately demonstrated efficient single protein expression from beads, DNA assembly on beads, and bead-based protein expression in microfluidic droplets, we next combined these approaches for the co-expression of multiple proteins within a single droplet, which has so far remained elusive. We concatenated three different genes by on-bead Golden Gate assembly as described before and encapsulated them in cell-free containing droplets. We confirmed by fluorescent confocal microscopy that multi-protein expression occurred in droplets containing only a single bead (Figure 4b). Compared to the monoscistronic expression (Figure 4a), multiscistronic expression led to an overall lower fluorescence signal. This was expected from previous co-expression bulk experiments (Figure 2) as limited expression resources need to be now shared by multiple templates. In summary, this experiment showed that multiple genetic programs can be easily assembled on beads and used as robust DNA templates for multi-protein expression.

## 4 Discussion

Cell-free reactions do not offer the same convenient phenotype to genotype link that many cell-based bioengineering methods rely on. In order to address this limitation, cell-free reactions can be encapsulated into droplets with a limiting dilution of DNA that distributes at most one DNA molecule per cell-like compartment [18, 21, 39, 40]. Unfortunately, the resulting DNA template concentration is typically too low for significant protein expression [39]. Several studies thus re-amplified a single DNA template within droplets before merging them with cell-free reactions [24, 41, 26]. However, microfluidic droplet merging or injection remains, in our opinion, the domain of specialized microfluidic labs and is not currently robust and easy enough for routine use.

We here instead immobilized high copy numbers of DNA templates on streptavidin-coated beads. Such beads are used for single-cell sequencing protocols and therefore widely available [42, 43]. We showed that in-droplet protein expression from a single DNA-loaded bead is similar to cell-free expression from DNA in solution. The problem was then how to get synthetic DNA onto beads such that sequences differ between but not within a bead. Various methods have already been developed: Kosuri and colleagues, for example, combined DNA barcoded beads and emulsion PCR into a highly multiplexed gene synthesis protocol for DNA fragments up to ≈ 700 bp in size [28, 29]. In another example, Lindenburg et al. built a more conventional site saturation DNA library of an affibody gene onto beads [30]. Their method consisted of chemical DNA immobilization followed by two different sets of restriction and ligation reactions. They then, in fact, screened this on-bead library in cell-free reaction droplets for enhanced target binding.

We were interested to find a simpler approach that would go beyond single-gene engineering and support the rapid testing of multi-gene programs. We developed two different methods for a solid-phase gene assembly. Both Golden Gate with a mix of BsaI and T4 ligase [36] or a single-enzyme DNA recombination with the recently discovered BxB1 integrase [37] allowed for efficient multi-gene assembly on beads. Golden Gate has the advantage of short ligation scars and all reagents are commercially available. BxB1 recombineering was yet more efficient to the point that we could not detect any significant difference in the expression level of the first and the last gene in the chain. Directly after gene assembly, we subjected beads to cell-free protein expression in droplet micro-reactors. We confirmed that a single bead per droplet was sufficient for the reliable co-expression of three different fluorescent proteins to easily detected levels.

The goal of our study was to develop a viable strategy that brings multi-gene expression into droplet emulsions but remains accessible to laboratories with little expertise in microfluidics. Consequently, we optimized our protocol on a commercially available microfluidic droplet generator and used off-the-shelf components and reagents throughout. Nevertheless, our study remains at a proof of concept stage. Further optimization and development are required for the implementation of an actual multi-gene *in-vitro* evolution application. One issue we identified stems from the tendency of micro-beads to clog up [35]. Such clusters lower the yield of single-bead droplets. They may also temporarily block microfluidic channels and perturb droplet composition leading to failures in protein expression. Furthermore, we did not explore the different options for screening for a particular phenotype. The best method for this task will depend on the application. Nonetheless, one clear option is fluorescent-assisted sorting. Cell-free water-in-oil droplets can be transformed into a double-emulsion using the same commercial microfluidics setup. FACS instruments can then easily be employed for fluorescent-activated droplet sorting [44, 45].

In conclusion, our combination of beads, cell-free system and microfluidic droplets offers a route towards potentially generating millions of reactions with diverse multi-gene DNA programs running in each droplet. Direct multi-gene assembly on beads moreover removes cloning as a major bottleneck in the design-build-test cycle while also maximising the number of DNA templates per drop. The whole platform can be readily implemented from commercially available instruments and reagents. We therefore hope that this methodology will open up the way for the microfluidics-based in-vitro evolution of more complex bio-engineered systems such as genetic circuits or metabolic pathways.

## 5 Supporting information

**supplement.pdf** supplementary material: figures (Fig.S1 - Fig.S5) and tables (Table S1 - Table S7)

**additional information** annotated DNA sequences and MATLAB scripts are available at https://github.com/strubelab/dropletXpress

## 6 acknowledgements

We thank David Conchouso for critical help with the setup of microfluidics as well as Daniela A. Garcia-Soriano and Marc Guëll for discussions and practical advice. We thank Lyazzat Bekish for purifying the LSSmOrange fluorescent protein. We are grateful to the KAUST Catalysis Center (KCC) and Ulrich Buttner, from the KAUST Nanofabrication Corelab, for providing materials and space for our microfluidic set-up. The KAUST Imaging and Characterization Core lab kindly provided training and assistance for confocal microscopy. This research was supported by the King Abdullah University of Science and Technology (KAUST) through the baseline fund and the Award No. URF/1/1976-21 from the Office of Sponsored Research (OSR).

## Notes

### Competing Interest Statement

The authors have declared no competing interest.

